# *In Vitro* Fertilization induces reproductive changes in male mouse offspring and has multigenerational effects

**DOI:** 10.1101/2024.11.06.622317

**Authors:** Eric A. Rhon-Calderon, Cassidy N. Hemphill, Alexandra J. Savage, Laren Riesche, Richard M. Schultz, Marisa S. Bartolomei

## Abstract

*In vitro* fertilization (IVF) is a non-coital method of conception used to treat human infertility. Although IVF is viewed as largely safe, it is associated with adverse outcomes in the fetus, placenta, and adult offspring life. Because studies focusing on the effect of IVF on the male reproductive system are limited, we used a mouse model to assess the morphological and molecular effects of IVF on male offspring. We evaluated three developmental stages: 18.5-day fetuses and 12- and 39-week-old adults. Regardless of age, we observed changes in testicular-to-body weight ratios, serum testosterone levels, testicular morphology, gene expression, and DNA methylation. Also, sperm showed changes in morphology and DNA methylation. To assess multigenerational phenotypes, we mated IVF and naturally conceived males with wild-type females. Offspring from IVF males exhibited decreased fetal weight-to-placental weight ratios and changes in placenta morphology regardless of sex. At 12-weeks-of-age, offspring showed higher body weights and differences in glucose, triglycerides, insulin, total cholesterol, HDL and LDL/VLDL levels. Both sexes showed changes in gene expression in liver, testes and ovaries, and decreased global DNA methylation. Collectively, our findings demonstrate that male IVF offspring exhibit abnormal testicular and sperm morphology and molecular alterations and transmit defects multigenerationally.

## Introduction

*In vitro* fertilization (IVF) is a widely used assisted reproductive technology (ART) to treat infertility (1). Since the first successful human IVF birth in 1978, this technology has dramatically evolved, resulting in improved pregnancy rates (2). Although considered safe, IVF pregnancies are associated with an increased risk of perinatal, neonatal, and placental complications, rare genetic syndromes, and possible long-term effects in the offspring in both humans and mice (3). One theory explaining adverse outcomes is that because IVF procedures occur during critical windows of extensive epigenetic reprogramming in gametes and preimplantation embryos (3, 4), external factors could impact normal development. In fact, embryo culture is a major contributor to adverse IVF outcomes (5). Although culture conditions used for IVF have been modified to improve success in fertilization rates and blastocyst formation (6), no concomitant improvement in offspring outcomes has been observed. Similarly, techniques such as vitrification and preimplantation genetic testing have been integrated into standard IVF cycles, aiming to enhance pregnancy success (3, 7). As IVF becomes more widely used due to factors like delayed childbearing, same-sex couples seeking parenthood, and an increasing incidence of infertility, understanding the potential adverse outcomes for the next generation becomes increasingly crucial (8). This concern is furthered by an interest in identifying multigenerational and transgenerational changes due to IVF and other ART procedures (9).

The mouse has emerged as an excellent model to study ART, simulating the human counterpart and enabling more invasive studies (10). Previous studies using mouse models have shown that IVF procedures are associated with a higher risk of metabolic syndromes in offspring, which includes alterations in glucose, insulin, triglycerides, cholesterol, and changes in the liver transcriptome and proteome compared to naturally conceived offspring (7, 11–19). Nevertheless, there is a paucity of information regarding potential effects on the reproductive system in ART-conceived offspring as well as the effects of traditional IVF on the next generation. Previous studies have shown that testicular or sperm maturation changes impact normal sperm function, which leads to adverse outcomes in sired offspring (20). A recent study using a mouse model demonstrated that metabolic changes in male IVF offspring, such as increased body weight, altered insulin levels and changes in fat deposition were transmitted to the second generation without the need for an additional insult, such as a high-fat diet. However, the reproductive system was not examined, and the mechanism underlying the multigenerational effects was not pursued (21).

Intracytoplasmic sperm injection (ICSI) is widely used for male specific infertility or when the fallopian tubes are obstructed (20). ICSI has similarities to conventional IVF, including embryo culture and transfer (22). Although men conceived via ICSI/IVF had similar sperm concentrations, total sperm counts, and total motile counts to those conceived naturally, subtle but significant differences were observed in sperm morphology and progressive motility, as well as in gonadotropin and serum testosterone levels (23). Another study, using mice, showed that male ICSI-conceived offspring not only showed changes in behavior and DNA methylation at promoter regions in testis DNA but also reported a transgenerational effect of ICSI, including changes in behavior and testicular levels of DNA methylation (9).

Although numerous adverse changes to placentas, liver and metabolic profiles have been observed in mice conceived by IVF (7, 11, 24, 25), to our knowledge, no study has focused specifically on the male reproductive system. To address this gap, we assessed the morphological and molecular effects of IVF on the testes and sperm of male offspring and then evaluated whether these adverse outcomes impacted the next generation. We report that IVF-conceived males exhibit changes in testicular-to-body weight ratios, decreased testosterone levels, alterations in seminiferous tubule morphology, DNA methylation perturbations in testes and sperm, and altered gene expression compared to naturally-conceived offspring. Moreover, second-generation offspring display metabolic perturbations and transcriptome changes in liver, testis and ovaries, and changes in global DNA methylation of testes and ovaries.

## Results

### Effects of IVF on testis weight and histology

To ascertain whether IVF could influence male reproductive function, we first assessed the effect of IVF on testes weight and histology using our mouse model (Figure 1A). Mean body and testicular weight at both 12 and 39 weeks were significantly higher in offspring of the IVF group compared to naturally-conceived controls (Naturals) (Figure 1B, C), which led to a significantly higher testicular-to-body weight ratio for the IVF group in 39-week-offspring (Figure 1D). Because testes are the primary producers of the key male sex hormone, testosterone, they play a critical role in both physical and reproductive health (26). Due to testosterone’s effect on multiple systems, abnormal testosterone levels can be indicators of underlying testicular and overall adverse health conditions (27). Accordingly, we determined whether an increase in testicular weight correlated with changes in testosterone levels and expression of the androgen receptor (AR) and found that both testosterone levels and AR expression in whole testis were significantly reduced in the IVF group at both 12 and 39 weeks compared to Naturals (Figure 1E).

**Figure 1.**
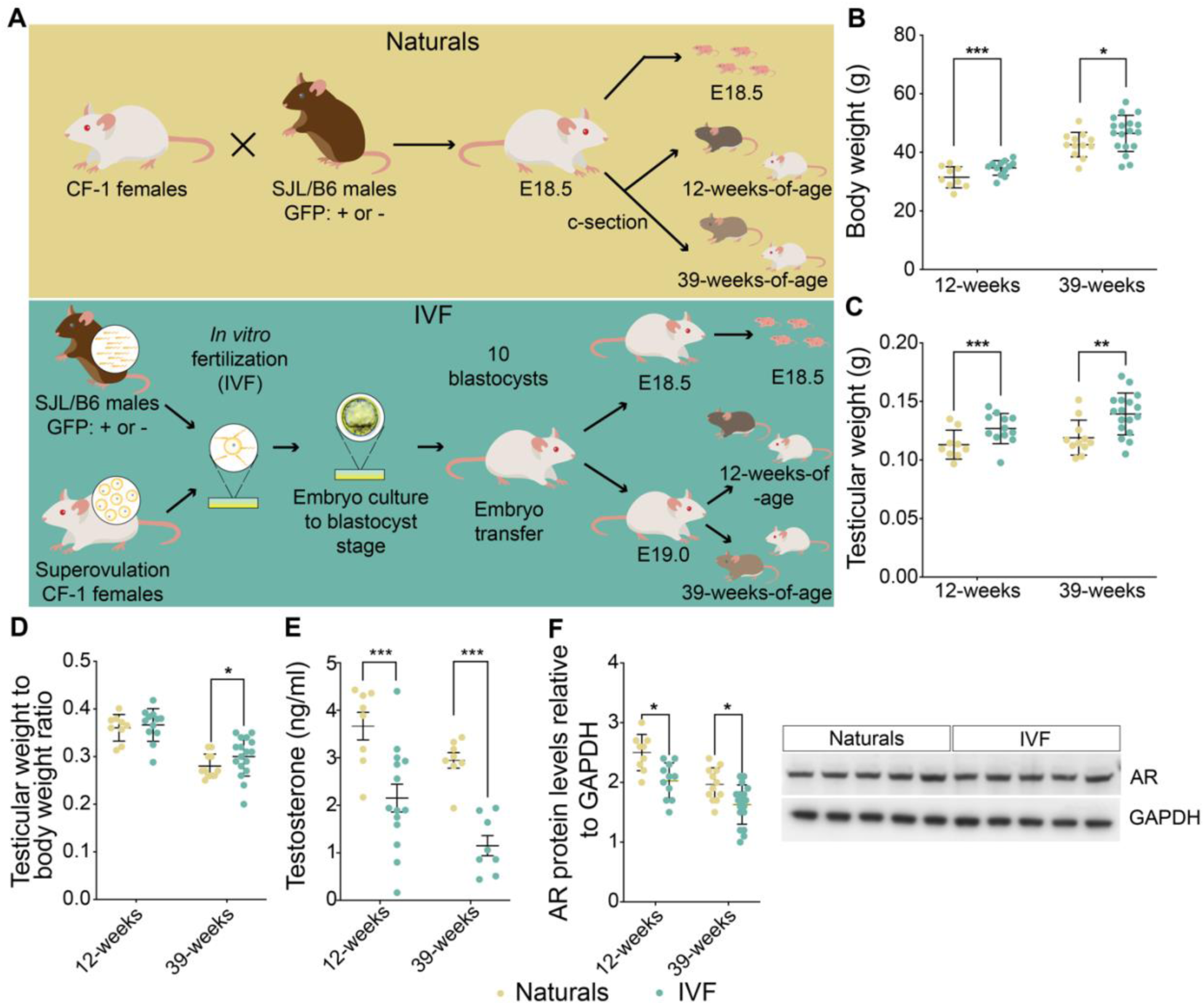
Mouse model and general parameters. (A) Mouse model: Top: Naturally conceived embryos (Naturals) were generated using SJL/B6 male mice and CF-1 female mice, and Bottom: IVF embryos were produced using capacitated sperm from SJL/B6 GFP+ or GFP- mice and eggs from superovulated CF-1 mice. Embryos were cultured to the blastocyst stage; 10 blastocysts were transferred to pseudopregnant females. At E18.5 pregnant females from both groups were c-sectioned and fetuses were delivered and collected for molecular analysis. For adult cohorts c-section-delivered pups at E18.5 (Naturals) or E19.0 (IVF) were fostered with wild-type dams and monitored until 12 or 39-weeks of age. Shown are: (B) Body weights before 6 h fasting, (C) testicular weights, (D) testicular to body weight ratios, (E) serum testosterone levels assayed by ELISA and (F) representative immunoblot testicular protein levels of AR relative to GAPDH assayed by westerns. Data are depicted as mean±s.e.m, *n*=10-12 per group. The black line represents the mean of each group. Statistical significance was determined by t-test, *P<0.05, **P<0.01 and ***P<0.001 when IVF groups compared to Naturals.

Spermatogenesis is a complex process involving the proliferation and differentiation of germ cells. Several factors could influence this process, including alterations in the seminiferous epithelium (28). Previous studies showed that seminiferous epithelium changes impair the proper elongation of spermatids, impact axoneme formation, and disturb interactions between Sertoli cells and germ cells that lead to impaired spermatogenesis (28). Seminiferous epithelium perturbations also arrest spermatogenesis at various developmental stages in mutant mice (29). Thus, any adverse changes to the seminiferous epithelium may lead to impaired spermatogenesis, impacting male fertility. We therefore histologically examined the seminiferous epithelium at 12 and 39 weeks (Figure 2A, B). Hematoxylin-eosin staining revealed an increase in seminiferous tubule cross-sectional area and the seminiferous epithelium per tubule at 39 weeks but not at 12 weeks (Figure 2C, F), whereas the height and area of the seminiferous epithelium were increased in both 12 and 39-week-offspring, (Figure 2D, E).

**Figure 2.**
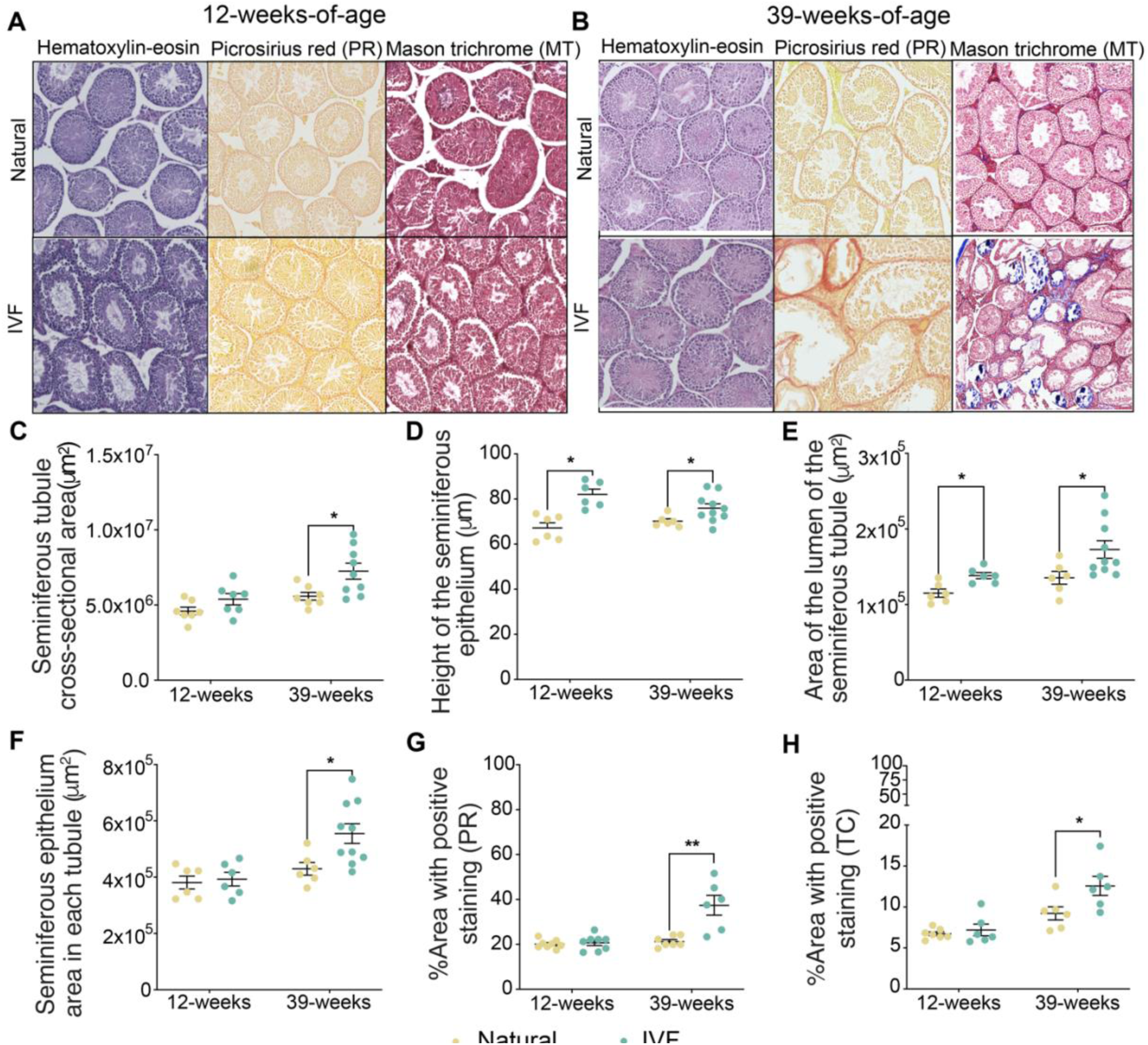
Testicular morphology analysis using hematoxylin-eosin, picrosirius red, and Masson’s trichrome staining. Testicular cross-sections at (A) 12-weeks-of-age, and (B) 39-weeks-of-age using hematoxylin-eosin, picrosirius red (PR) and Masson’s trichrome (Magnification is 4x). Morphology of the seminiferous tubules in hematoxylin-eosin slides, including: (C) Whole cross-sectional area, (D) Height, (E) Lumen area, and (F) Total seminiferous epithelium area. Percentage of positively stained area with PR (G) and Masson’s trichrome (H) is depicted. The black line represents the mean of each group. Each data point represents an individual conceptus from a minimum of five different litters (n=5-10/group). Data are expressed as mean±s.e.m. Statistical analysis between groups was done by t-test and * represents a significant difference of *P*<0.05, when comparing IVF against Naturals.

In testes, collagen, a major component of the testicular basement membrane, is located at the base of the seminiferous epithelium (30). The basement membrane has multiple functions, including germ cell movement and communication between different cell types (31). Disruption of this matrix is associated with azoospermia (32). Additionally, excessive collagen is associated with an increase in tunica albuginea weight and thickening of the basement membrane, as seen in fibrosis and aging models (30). We therefore assessed the effect of ART on the extracellular matrix in testis, specifically the collagen density, using two staining protocols: picrosirius red (PR), which quickly and efficiently assesses collagen fiber content, organization, and orientation (33) and Masson’s trichrome (MT), which not only detects the extent of collagen fiber deposition and but also can distinguish muscle fibers (red), collagen (blue), and nuclei (black) simultaneously (34). Both methods revealed increased staining intensity exclusively at 39 weeks (Figure 2G, H). To validate this increased staining, we assayed *Col1a1* RNA and protein by RT-qPCR and immunoblotting, respectively, as well as another extracellular matrix marker, vascular cell adhesion protein 1 (*Vcam1*), all of which were increased in IVF offspring compared to Naturals (Supplemental Figure 1). Taken together, these results indicate an increased thickening of the tubular walls and a greater degree of disorganization in tubules with higher collagen content in our IVF male offspring. Additionally, this increase in collagen deposition contributed to the increased testicular weight in IVF offspring.

### Effect of IVF on sperm count, morphology, and DNA methylation

Given the IVF-associated changes in testis morphology, we next examined the effect of IVF on several sperm parameters. Although there was no significant difference in sperm counts from 12-week-old IVF-offspring, there was greater variability in sperm number (Figure 3A). When testes size and sperm count were correlated, IVF offspring with larger testes had lower sperm counts (Supplemental Figure 2), and the percentage of sperm with abnormal morphology was increased (Figure 3B). No data are available for the 39-week time point because sperm were immediately frozen, and no subsequent characterization was performed.

**Figure 3.**
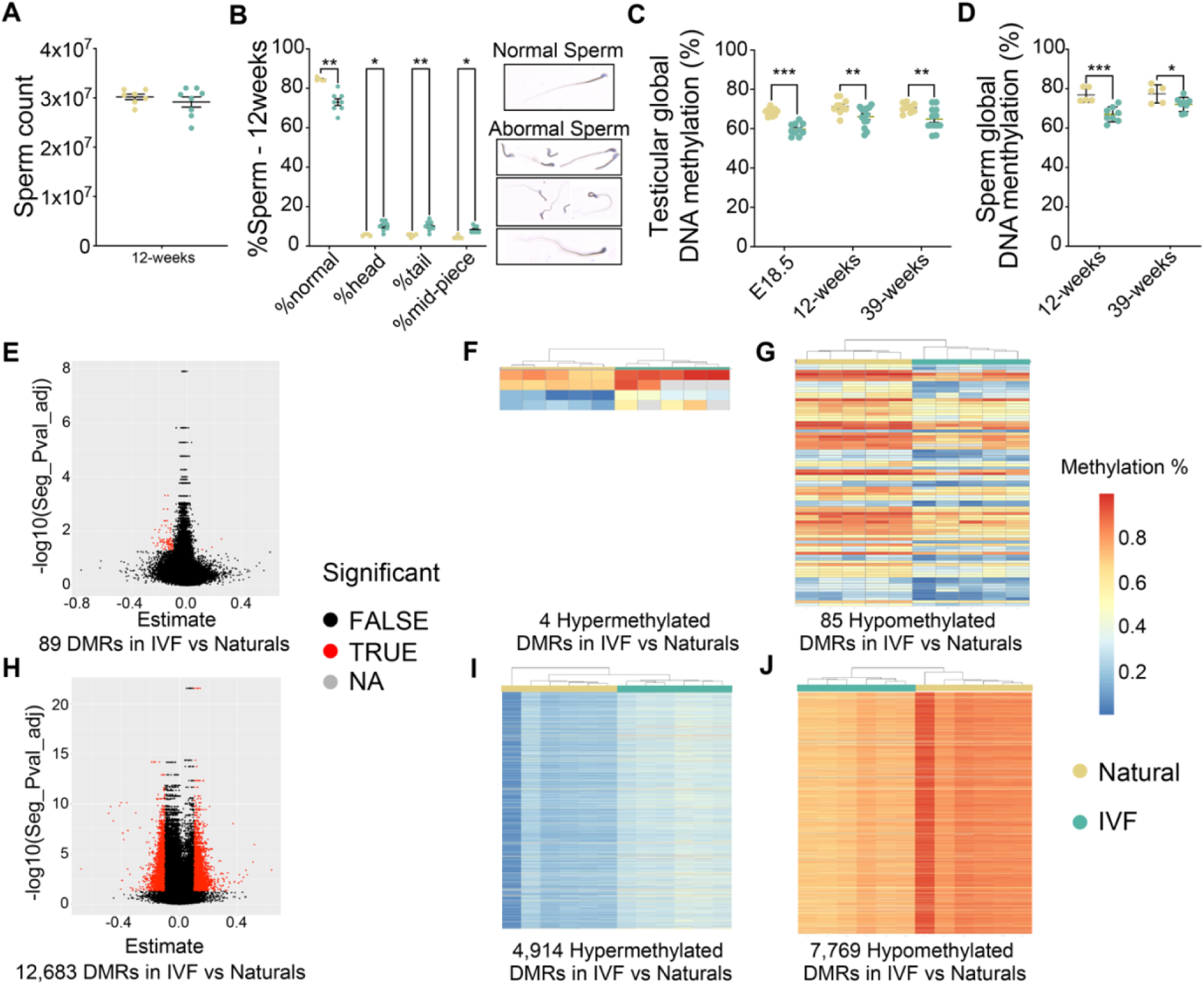
Sperm parameters and testicular DNA methylation. (A) Sperm count using a brightfield microscope, (B) Sperm morphology quantification using eosin 1% staining where sperm is categorized as normal (%normal), abnormal head (%head), abnormal tail (%tail) and abnormal mid-piece (%mid-piece), (C) Testicular global DNA methylation and (D) Sperm global DNA methylation using LUMA for 12 and 39-week Natural and IVF offspring. Infinium Mouse Methylation BeadChip sperm analysis including, (E) volcano plot for DMRs at 12-weeks, 12-weeks Heatmap with (F) hypermethylated DMRs and (G) hypomethylated DMRs, (H) volcano plot for DMRs at 39-weeks, 39-weeks Heatmap with (I) hypermethylated DMRs and (J) Hypomethylated DMRs. Each data point represents an individual conceptus from a minimum of five different litters (n=10-15/group). Data are expressed as mean±s.e.m. Statistical analysis between groups was done by t-test and *, **, *** represent a significant differences of *P*<0.05, *P*<0.01 and *P*<0.001, respectively, when comparing IVF to Naturals.

IVF is also associated with changes in whole-genome average DNA methylation (7, 11, 24, 25). We therefore assessed global DNA methylation using a luminometric methylation assay (LUMA) on testis DNA at E18.5, 12 and 39 weeks, and sperm DNA at 12 and 39 weeks. We observed statistically significant decreased DNA methylation in testes and sperm from IVF-conceived male offspring compared to Naturals at all timepoints. (Figure 3C, D). To identify specific regions with altered DNA methylation that could potentially be transmitted to offspring, we examined DNA methylation in sperm from 12- and 39-week-old IVF-conceived mice using the Infinium Mouse Methylation-12v1-0 BeadChip array, which assays approximately 285,000 CpG probes distributed across the genome (7, 24, 35, 36). As previously described (7, 35), we first used all probes to determine the differentially methylated loci (DMLs) that had a 0.05 difference between groups. With these DMLs we then determined differentially methylated regions (DMRs) that are associated with a differentially methylated probe on the array. We considered a DMR as a genomic region that had a 10% difference in methylation between groups. A 10% difference is considered the threshold for DNA methylation to influence gene expression. Heatmap and PCA analysis by probe passing rate, group and age, showed a differential clustering between sperm from IVF and Natural offspring (Supplemental Figure 2). At 12-weeks, we observed 89 DMRs, 4 hypermethylated DMRs and 85 hypomethylated in IVF compared to Naturals (Figure 3E). Gene ontology analysis failed to detect significant association with specific biological functions at 12 weeks. Finally, in 39-week sperm samples, we observed that IVF-conceived males exhibited 12,683 DMRs, 4,914 hypermethylated DMRs and 7,769 hypomethylated compared to Naturals (Figure 3F). Gene ontology analysis showed that the most affected pathways were related to cell-cell interactions, retrotransposon silencing, regulation of membrane potential, signal transduction, cell morphogenesis and regulation of actin filament processes (Figure 3G and 3H). Together these results show that IVF procedures impacted both testicular and sperm DNA methylation with possible effects on the next generation, and that age increased the number of DMRs in sperm.

### Effect of IVF on testicular gene expression

To ascertain whether the changes in DNA methylation are associated with changes in gene expression of IVF-conceived males, we performed RNA-seq on fetal and adult testes collected at E18.5, 12 weeks, and 39 weeks. PCA showed a clear separation between IVF and Naturals (Supplemental Figure 3A-C). Volcano plots revealed 57 upregulated and 78 downregulated genes at E18.5 (Figure 4A), 17 upregulated and 57 downregulated genes at 12 weeks (Figure 4B), and 64 upregulated and 29 downregulated genes at 39 weeks (Figure 4C). Not all the DEGs at 12 weeks were observed at 39 weeks. Testes from IVF-conceived males at E18.5 had the greatest number of DEGs, and this difference did not appear to worsen with age. The heatmap for all DEGs showed tight clustering for all Naturals, with IVF groups clustering together (Figure 4D-F). Gene ontology analysis showed dysregulation in transmembrane transportation and testicular function at E18.5 but failed to detect significant association with specific biological functions at 12 and 39 weeks (Figure 4G). Finally, we determined if the DEGs in testis overlap with affected sperm DMRs. At 12 weeks we did not observe any overlap between the DEGs and DMRs, while at 39 weeks we observed that one downregulated gene: *Ttc22*, and 8 upregulated genes: *1700019P21Rik*, *Cd24a*, *Gm31447*, *Nedd9*, *8430426J06Rik*, *Dazl*, *Sycp1* and *Adamts9* expression was linked with hypermethylated DMRs or hypomethylated DMRs respectively. (Supplemental Figure 3E and 3F). This data suggests little correlation between changes in gene expression and DNA methylation, indicating that the observed changes in gene expression may be due to other epigenetic modifications not measured in this study.

**Figure 4.**
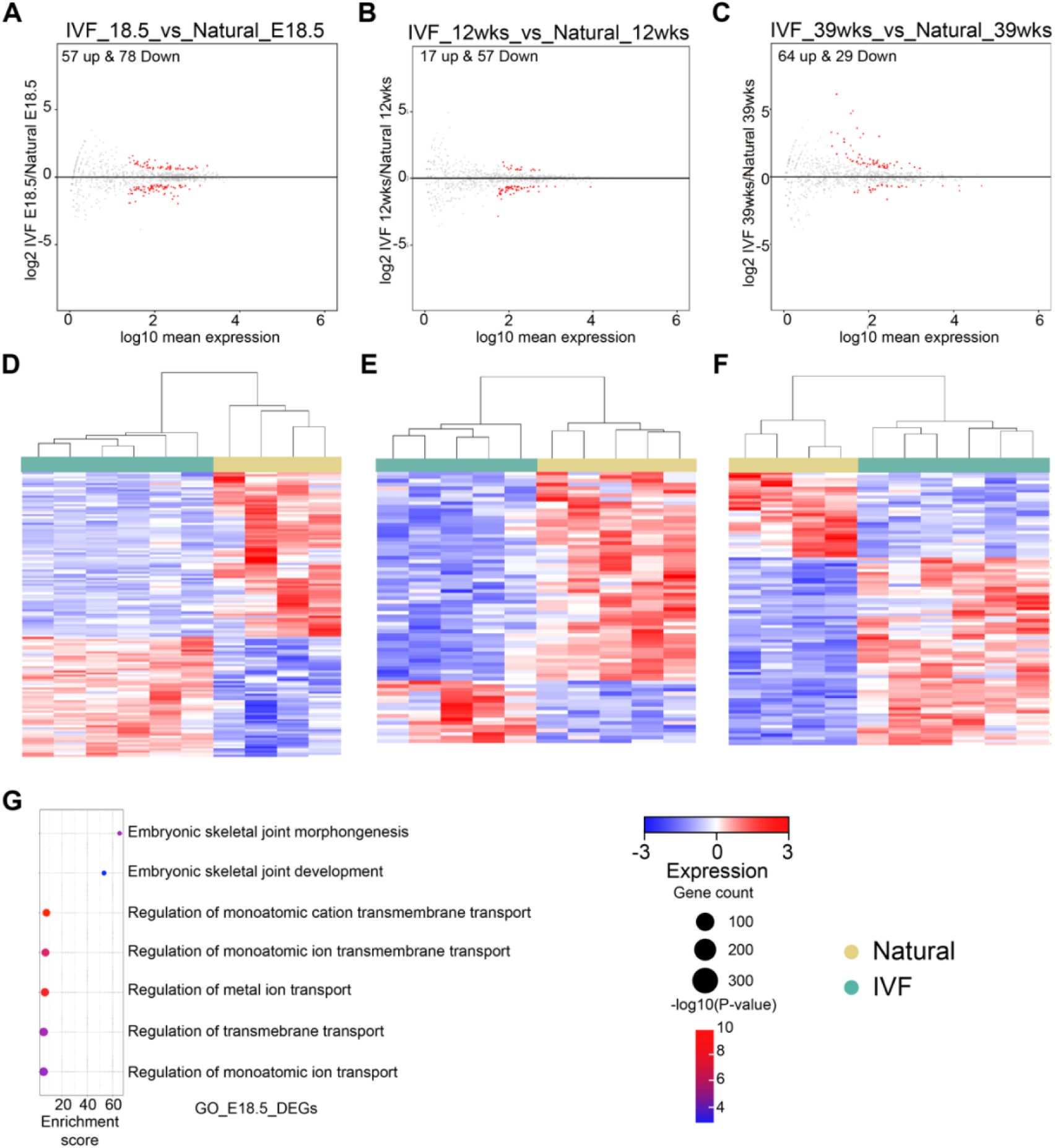
Transcriptome analysis of testicular tissues by RNA-seq. Volcano plots showing DEGs for (A) E18.5 gonads and testis at (B) 12-weeks, and (C) 39-weeks IVF vs Natural. Heatmap for log2-transformed expression levels obtained from RNA-Seq of DEGs in (D) E18.5 gonads, testis at (E) 12-weeks and (F) 39-weeks. Gene ontology analysis for relevant affected pathways: (G) E18.5 testis. Color boxes at the top denote the experimental group.

### Multigenerational Effects of IVF on placental to fetal weight ratios in F2 concepti

Considering that sperm DNA methylation was perturbed in IVF-conceived males, we determined whether these changes could be transmitted to their offspring. Accordingly, 12-week-old IVF-conceived males and Naturals were mated with CF-1 wild-type females, and concepti (F2) were isolated at E18.5 (Figure 5A). Litters sired by IVF-conceived males had fewer live pups compared to those sired by Naturals (Figure 5B). Because we previously identified sex differences in fetal and placental weight in the first generation (7, 36), we performed all analyses by sex. In F2 males, although the mean placental weight was unchanged at E18.5 (Figure 5C), the mean fetal weight was decreased for IVF concepti compared to Naturals (Figure 5D). In F2 females, both mean placental weight and mean fetal weight were decreased for IVF concepti compared to Naturals at E18.5 (Figures 5E and 5F). The reduction in fetal weight, with or without a decrease in placental weight, resulted in significantly reduced fetal:placental weight ratios for F2 IVF compared to Naturals (Figure 5G, 5H). Finally, to address if IVF has a multigenerational impact on the area of the junctional and labyrinth regions in the placenta, as previously observed on IVF offspring (24, 25), we used hematoxylin-eosin staining at E18.5 (Supplemental Figure 4A-F). The percentage of labyrinth and junctional zone area was significantly lower and higher, respectively, in F2 IVF placentas compared with Naturals (Supplemental Figure 4B, 4C, 4E and 4F) in both female and male offspring as previously observed in IVF fetal offspring (7).

**Figure 5.**
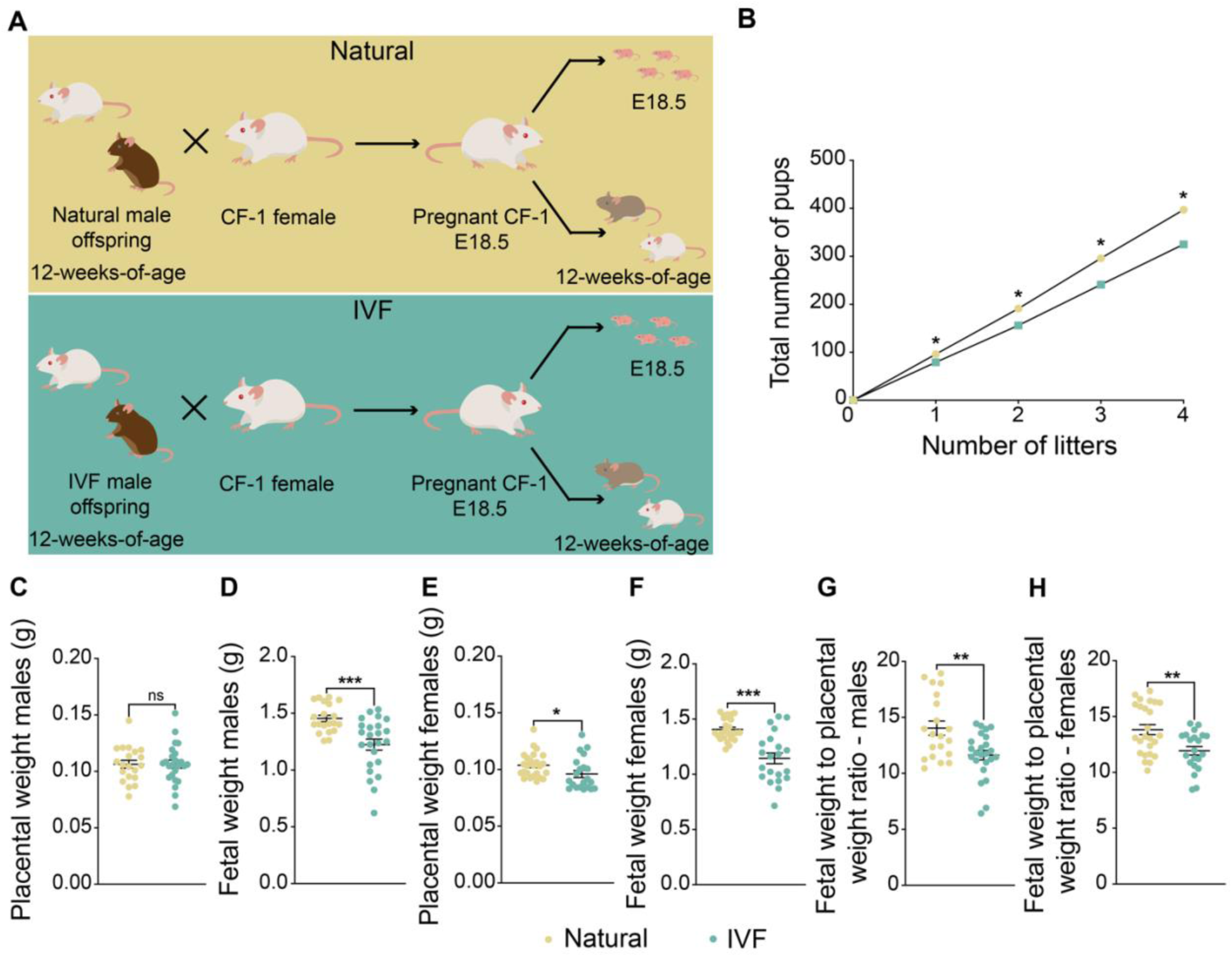
Multigenerational impacts of IVF. (A) Mouse model: Top: F2 Naturally conceived offspring were generated using Natural male offspring and CF-1 female mice, and Bottom: F2 IVF offspring were produced by using IVF male offspring and CF-1 female mice. E18.5 pregnant females from both groups were c-sectioned and concepti were delivered and collected for molecular and histological analysis. For adult cohorts naturally-delivered pups were maintained until 12-weeks-of-age. (B) Total number of pups is depicted for 4 consecutive litters, with data presented as summary of total pups after each litter from 4 breeding pairs/group. Placental (C) and fetal (D) weight for male offspring, and placental (E) and fetal (F) weight for female offspring are shown. The fetal weight to placental weight ratio for male (G) and female (H) offspring are depicted. Data are mean±s.e.m, *n*=16-20 per group. Black lines represent the mean of each group. Statistical significance was determined by t-test, *P<0.05, **P<0.01 and ***P<0.001, when IVF groups are compared to Naturals.

### Effect of IVF on male F2 adult offspring metabolism

We and others previously determined that metabolism is perturbed in IVF-conceived offspring (7, 11, 13, 18, 19). To determine whether the F2 progeny sired by IVF-conceived males likewise exhibit adverse health outcomes, we measured the body weight of a cohort of F2 male offspring until 12-weeks-of-age (Figure 5A), and subsequently euthanized the mice for metabolic analyses. Male F2 offspring sired by IVF-conceived males had higher body weight compared to F2 offspring sired by Naturals (Figure 6A). Serum glucose, insulin, total triglycerides and LDL/VLDL were also elevated in F2 offspring relative to Naturals (Figure 6B-E). Total cholesterol was decreased, which was driven by reduced HLD (Figure 6E). We also recorded organ weights and calculated their ratio to body weight. F2 offspring sired by IVF-conceived males exhibited decreased brain and testis weight relative to body weight compared to F2 offspring sired by Naturals (Figure 6F), whereas pancreas and kidney weights relative to body weight were increased (Figure 6F). Similar phenotypes were also observed in the original IVF male offspring (7, 11).

**Figure 6.**
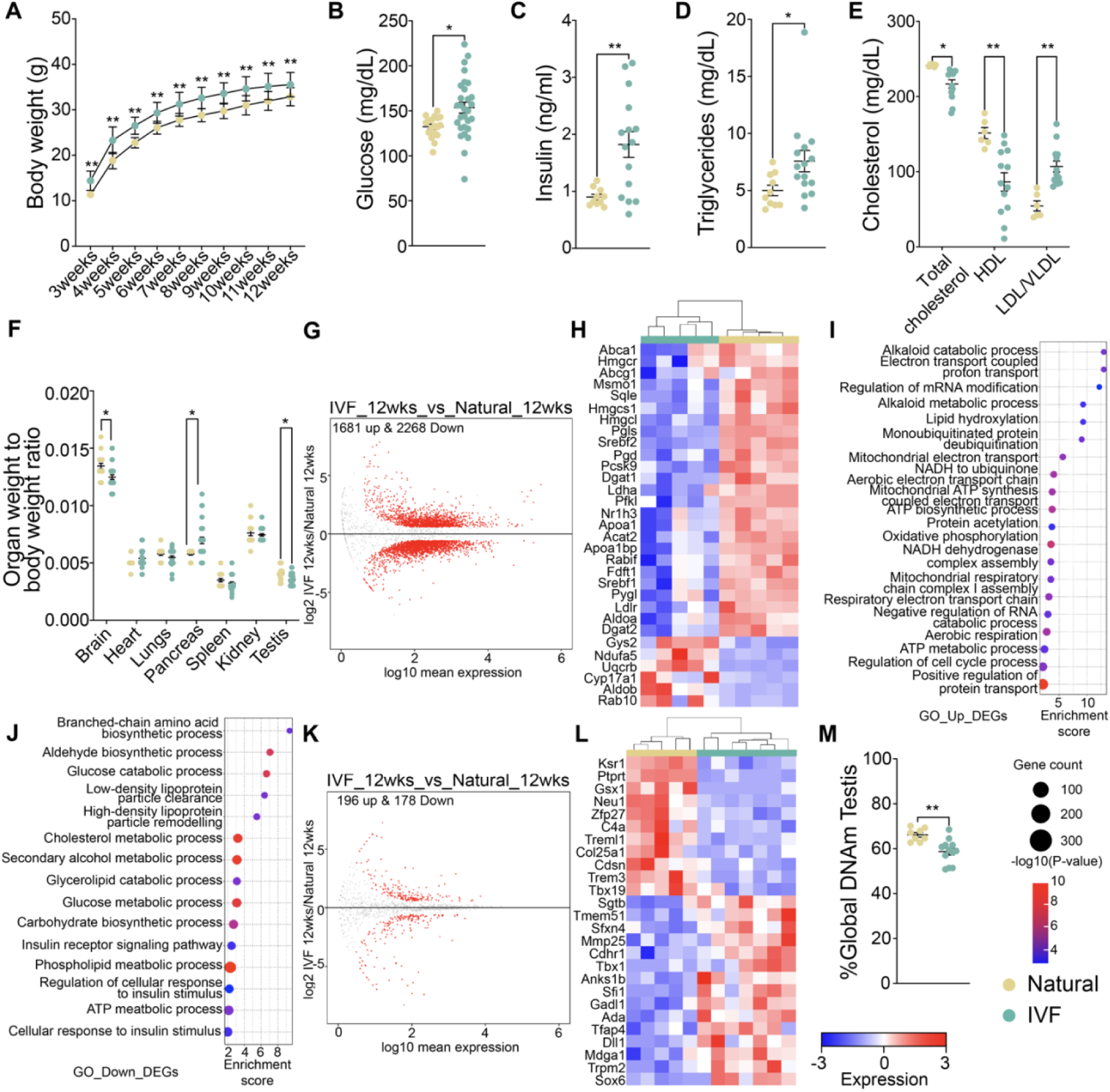
Multigenerational impact of IVF on male offspring. (A) Body weights were taken from 3-12-weeks-of-age. Metabolic screening included (B) glucose, (C) insulin, (D) triglycerides and (E) total cholesterol, HDL and LDL/VLDL. (F) The organ weight to body weight ratio is shown. (G) Volcano plot showing DEGs for liver RNA-seq, (H) Heatmap for log2-transformed expression levels obtained from RNA-seq of DEGs involved in cholesterol, triglycerides, insulin and glucose metabolic pathways. Gene ontology analysis for pathways involved in metabolism using (I) upregulated DEGs and (J) downregulated DEGs. (K) Volcano plot showing DEGs for testis RNA-seq, (L) Heatmap for log2-transformed expression levels obtained from RNA-seq of DEGs involved in spermatogenesis and testis differentiation, and (M) Global DNA methylation of F2 testis at 12-weeks by LUMA. Data are shown as mean±s.e.m, *n*=10-15 per group. The black line represents the mean of each group. Statistical significance was determined by t-test, *P<0.05 and **P<0.01, when IVF compared to Natural. For RNA-seq, color boxes at the top denote the experimental group.

In a previous study, we showed that 12-week IVF-offspring exhibited changes in liver gene expression (11). We performed RNA-seq on a subset of liver and testis samples from male F2 progeny. For liver, PCA showed a clear separation between the IVF and Natural groups (Supplemental Figure 5A), with 1,486 upregulated and 1,936 downregulated genes observed when comparing IVF and Natural F2 offspring (Figure 6G). A subset of DEGs associated with metabolic pathways is shown in a heatmap (Figure 6H). Gene ontology analysis revealed dysregulation of pathways involved in cholesterol and phospholipid metabolism, glucose and insulin processes (Figure 6I, J). These results are consistent with the metabolic panel and demonstrated that the changes in metabolic measurements are the result of changes in the liver transcriptome. Additionally, we found that some of our liver DEGs, such as *Irs1*, *Irs2*, *Ripk2*, *Rspo1* and DNA JC genes, have been previously associated with insulin and glucose resistance (37), which would explain the higher insulin and glucose levels revealed in the metabolic panel. Additionally, we tested if there was a normal correlation between glucose and insulin, while Naturals show a positive correlation IVF show a more variable trend and not correlated the one with the other as expected in a metabolic syndrome (Supplementary Figure 8).

To determine whether DEGs observed in livers of F1 and F2 were similar, we conducted an overlap analysis using our previously obtained data (11). F2 offspring had more DEGs compared to F1 (3,949 vs 17), with no overlapping genes. Although the difference in gene expression could reflect the biology of the IVF males, in this study we doubled the number of samples analyzed, which provided more power to detect differences between the two groups. Nevertheless, considering the pathway analysis performed in our previous study (11), cholesterol and triglycerides pathways are equally affected in both F1 and F2 offspring.

For testis, PCA showed a clear separation between F2 offspring from IVF and Natural groups (Supplemental Figure 5B) and volcano plots comparing F2 offspring revealed 196 upregulated and 178 downregulated genes (Figure 6K). Some of the most affected genes involved in testicular function and development are shown in a heatmap with differential clustering between groups (Figure 6K). Gene ontology failed to detect significant association with specific biological functions.

To determine whether the changes in testis DNA methylation persisted into the next generation, we analyzed global DNA methylation in the testis of F2 offspring. Like the male IVF offspring, total global testicular DNA methylation was decreased in F2 IVF offspring compared to the natural conception group (Figure 6M). Taken together, these findings demonstrate that the changes observed in male IVF offspring have a multigenerational and persistent negative impact on metabolism, DNA methylation, and gene expression in the liver and gonads of F2 male offspring.

### Effect of IVF on female F2 adult offspring metabolism

Given the sex differences in outcomes in ART-conceived mice (7, 11, 18), we conducted a similar set of experiments to those described above. Female offspring from IVF-conceived males had a higher body weight compared to offspring sired by Naturals (Figure 7A). Serum glucose was decreased, whereas insulin, and total triglycerides were increased and displayed a greater variability in the F2 IVF females compared to Naturals (Figure 7B-D). In contrast, cholesterol, although not statistically different, displayed greater variability, and HDL and LDL/VLDL were decreased in F2 IVF offspring compared to Naturals (Figure 7E). Further, F2 offspring sired by IVF-conceived males showed a decrease in ovarian weight relative to body weight compared to Naturals (Figure 7F), and, as in males, pancreas and kidney weights relative to body weight were increased (Figure 7F). Most of these changes were previously observed in female IVF offspring (7, 11). As in males, we also tested if there was a normal correlation between glucose and insulin (Supplementary Figure 8).

**Figure 7.**
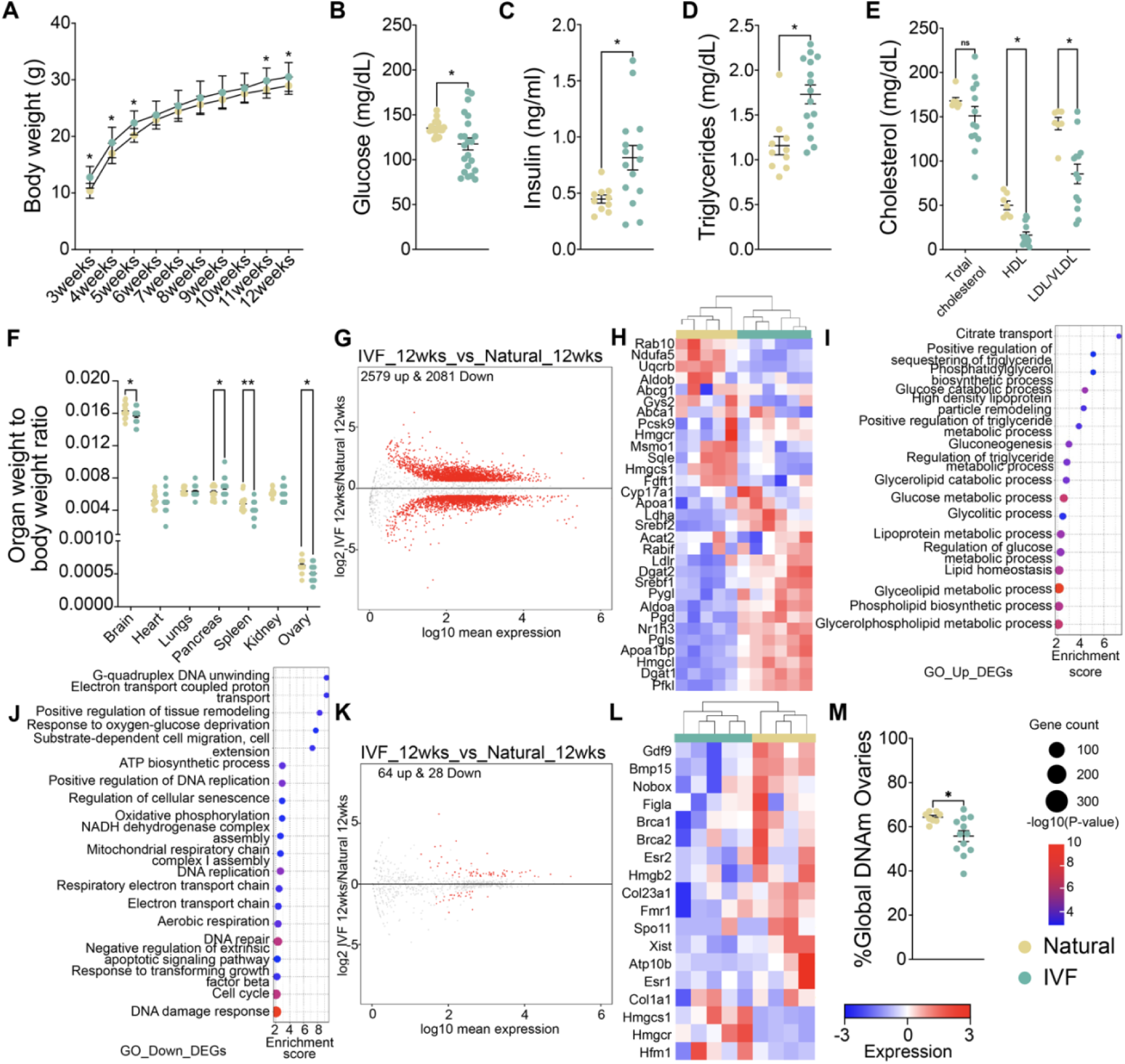
Multigenerational impact of IVF on female offspring. (A) Body weights were taken from 3-12-weeks-of-age. Metabolic screening included (B) glucose, (C) insulin, (D) triglycerides and (E) total cholesterol, HDL and LDL/VLDL. (F) The organ weight to body weight ratio is shown. (G) Volcano plot showing DEGs for liver RNA-seq, (H) Heatmap for log2-transformed expression levels obtained from RNA-seq of DEGs involved in cholesterol, triglycerides insulin and glucose metabolic pathways. Gene ontology analysis for pathways involved in metabolism using (I) upregulated DEGs and (J) downregulated DEGs. (K) Volcano plot showing DEGs for ovary RNA-seq, (L) Heatmap for log2-transformed expression levels obtained from RNA-seq of DEGs involved in ovarian function, and (M) Global DNA methylation of F2 ovary at 12-weeks by LUMA. Data is depicted as mean±s.e.m, *n*=10-15 per group. The black line represents the mean of each group. Statistical significance was determined by t-test, *P<0.05, and **P<0.01 when IVF compared to Natural. For RNA-seq color boxes at the top denote the experimental group.

We also performed RNA-seq on a subset of liver and ovary samples to determine whether gene expression was perturbed in female F2 progeny sired by IVF-conceived males. Similar to male offspring, PCA showed a clear separation between the IVF and Natural groups (Supplemental Figure 6A), and volcano plots revealed 2,579 upregulated and 2,081 downregulated genes (Figure 7G). A heatmap depicts DEGs involved in cholesterol, triglycerides, insulin and glucose metabolism (Figure 7H). Gene ontology analysis for upregulated (Figure 7I) and downregulated DEGs (Figure 7J) revealed that the most affected pathways are involved in glucose, cholesterol, lipid, and energy pathways, correlating to the observed changes in the metabolic panel. As in males, these results are consistent with the metabolic panel and demonstrated that the changes in metabolic measurements are the result of changes in the liver transcriptome. Although DEGs in males were associated with insulin resistance, DEGs in females were associated with diabetic syndrome.

When RNA-seq data from ovary were analyzed, PCA again showed a clear separation between F2 offspring sired by IVF-conceived males and those conceived by Natural males (Supplemental Figure 6B), with volcano plots showing 64 upregulated and 28 downregulated genes (Figure 7K). As with liver, we selected a subset of DEGs involved in ovarian function and displayed them on a heatmap (Figure 7L). GO analysis revealed that ovarian metabolism and hormone production were affected (Supplemental Figure 6C). As in males, we determined if DEGs observed in F1 and F2 offspring livers were similar. Again, F2 showed higher number of DEGs compared to F1 (4,660 vs 12), and none of the genes overlap (Supplemental Figure 6). Pathway analysis performed on our previous work (11) showed that cholesterol and triglycerides pathways are affected in F1 and the same pathways were affected in our F2 offspring.

To determine whether the DEGs observed in both the liver and gonads were similar between females and males, we conducted an overlap analysis. We found that 2,843 DEGs were shared in the liver, mostly involved in glucose, cholesterol, and triglyceride pathways, consistent with the metabolism panel. Additionally, we identified 1,817 DEGs unique to females and 1,106 unique to males (Supplemental Figure 7A). In gonads, 2 DEGs were shared, with 88 unique to females and 374 to males (Supplemental Figure 7B). Finally, as in males, we assessed whether changes in male IVF offspring could impact global DNA methylation in F2 female gonads. Global DNA methylation in the ovary was decreased in F2 IVF offspring compared to Naturals (Figure 7M). These ovarian changes suggest a multigenerational effect.

## Discussion

### IVF negatively impacts testicular and sperm parameters compared to Naturals

IVF is often considered an environmental exposure due to the *in vitro* manipulation of gametes and preimplantation embryos when dramatic epigenetic reprogramming occurs (38). A likely outcome of altered epigenetic reprogramming during these early stages of development is that multiple tissues would be affected, e.g., fetus, placenta, and liver (7, 11, 13, 18, 19). The results reported here significantly extend the number of affected tissues by demonstrating that IVF negatively impacts male reproductive health, noting that the reproductive system is extremely sensitive to environmental exposures, with potential consequences for both reproductive and multigenerational health (39).

Male offspring conceived via IVF exhibit significantly decreased levels of testosterone and androgen receptor. Testosterone is a crucial male hormone, essential for maintaining reproductive health and overall physiological functions (26, 27). Previous studies in models of obesity and metabolic syndrome have shown that males with reduced testosterone levels also exhibit a decline in AR expression due to a positive feedback loop (40). Although these hormonal changes may correlate with previously characterized metabolic syndromes, it is possible that changes during preimplantation and post implantation development directly influence gene expression pathways involved in testosterone production and AR regulation. Alterations in both the number and functional capacity of the cells responsible for testosterone production could also be a contributing factor in reduced AR expression.

Because of differences in testes weight and testosterone, we interrogated the seminiferous tubules of IVF male offspring, identifying alterations in the seminiferous epithelium. Studies in mice, rats, and humans have shown that alterations in the seminiferous tubules can impair spermatogenesis, affecting key processes such as spermatid elongation, axoneme formation, and interactions between Sertoli cells and germ cells (28). Depending on the severity of the disruption, these changes can also impact the timing and overall efficiency of spermatogenesis (28). Most environmental exposures and metabolic syndromes are associated with an increase in tubule area; however, in our study, we observe not only an increase in overall tubule area but also expansion of the epithelial membrane that may reflect dysregulated spermatogenesis and cell division.

Another potential cause of the increase in both testicular weight and overall tubule area is the over-deposition of collagen and laminin, which is typically seen in fibrosis of the testis (41). At 39 weeks, we observe higher collagen deposition in the seminiferous epithelium membrane, which could disrupt normal cell distribution and lead to the tubule overgrowth observed in IVF offspring. Excessive collagen deposition is linked to various pathologies, including fibrosis, Klinefelter syndrome, inflammation, infection, aging, and physical stress (41, 42). Our results are consistent with fibrosis and early aging. Future studies will address whether this phenotype is associated with specific gene mutations or changes in gene expression related to collagen regulation and the maintenance of the epithelial membrane.

Changes in testosterone levels and morphology provide a plausible explanation for the negative impact on spermatogenesis. Consistently, RNA-seq on testicular tissues identified dysregulation in genes involved in hormone production and regulation, as well as extracellular matrix composition and maintenance. Among the dysregulated genes in the testis, several were related to collagen production, which reflects the reported phenotype. Although in our model the collagen and morphology changes are explained by the dysregulation of genes important for these pathways, we also hypothesized that some of these changes could represent a response to the metabolic syndrome affecting the IVF male offspring (43). Previous studies in models of obesity and alcohol exposure have demonstrated that progeny of exposed animals exhibit gene expression dysregulation across multiple systems, including the testis. These changes have been associated with inherited phenotypes driven by altered sperm maturation due to disruptions in overall spermatogenesis (43–46).

Although the impact of IVF on sperm in IVF-conceived males could be due to abnormal seminiferous tubules, metabolic syndromes and environmental exposures have an impact on the number, morphology and molecular integrity of the sperm (47, 48). In humans, IVF male offspring did not present with changes in sperm count but did have an increase in sperm abnormalities (23). Here we also did not observe differences in sperm number at 12 weeks, but we do observe altered morphology, with most of the changes related to head and acrosome abnormalities. Ultimately, an increase in the number of sperm abnormalities will impact the success of fertilization and compromise male fertility (49).

Because of previously identified changes in DNA methylation in IVF offspring reported by our group and others (7, 13, 24, 25), we examined DNA methylation in both testis and sperm. Both testicular and sperm DNA methylation were globally impacted, which likely results from alterations in normal reprogramming during embryo formation. We also analyzed the sperm DNA methylation landscape in greater depth to determine the possibility of transmitting an abnormal epigenome to the next generation. A normal epigenetic landscape in sperm includes DNA methylation marks, retained histones and protamines, and non-coding RNAs (50). At both 12 and 39 weeks, we observed changes in DNA methylation, with a greater number of affected regions at the later age. Because environmental exposures can alter the DNA methylation patterns of sperm, negatively impacting the next generation (50, 51), IVF may induce similar effects, compromising sperm integrity and contributing to the multigenerational changes reported here. Interestingly, we did not observe a specific genomic feature affected, instead all genomic regions were affected as we have seen before in previous studies in other tissues (5, 7).

### IVF has a multigenerational negative impact

To our knowledge few studies have focused on the F2-generation of IVF offspring. Previous studies in mice reported that offspring from IVF/ICSI conceived-males exhibited altered fasting glucose and insulin levels (17, 21), with more pronounced changes when the offspring were exposed to a high-caloric diet (17, 21, 52). Because male gametes from IVF-conceived males have defects, e.g., changes in DNA methylation, we assessed whether such changes are observed in F2 offspring from IVF sires. At term (E18.5), we observed that sired offspring exhibited lower fetal and placental weights, which affected the fetal-to-placental weight ratio. Changes in this ratio are associated with altered placental efficiency (53) and are consistent with the small for gestational age (SGA) classification; SGA is linked to multiple paternal factors, including age, obesity, and other metabolic syndromes (54). Thus, the changes we observe in the F2 generation might be a consequence of the metabolic outcomes caused by IVF in the first generation, which are transmitted via affected sperm. Finally, we followed F2 mice to 12-weeks-of-age — a time point at which we have previously identified significant changes in metabolic phenotypes in both female and male IVF offspring (7, 11). We observed that whereas most individuals appeared healthy, some exhibited altered metabolism, which mirrored the changes observed in the F1 generation (7, 11). These metabolic changes reflect those previously associated with metabolic syndrome in both humans and mice (13, 18, 55). Interestingly, we observed sex specific adverse outcomes in the F2 from IVF offspring, including a higher risk of insulin and glucose resistance in males, and a diabetic phenotype in females. These sex-specific differences can arise either by sex-linked genes and/or hormones.

The changes observed in our model are likely the result of the cumulative effects of ART, such as the longer culture to reach blastocyst stage, and the metabolic syndromes developed by the IVF offspring fathers. Currently, no human studies have focused on the multigenerational effects of IVF largely due to the age of this population. The mouse model is advantageous for these studies because mice are sexually mature at 6-8 weeks, samples are easily accessible, external factors can be controlled, and the specific effects of ART interventions can be identified. Our findings highlight the importance of monitoring the long-term health of IVF offspring and subsequent generations.

In summary (Figure 8), we demonstrate that IVF offspring exhibit molecular and morphological changes in testis and sperm that affect not only the reproductive system but also other systemic functions. Additionally, these changes have a significant impact on the progeny of IVF offspring, leading to an increased risk of developing metabolic syndromes in adulthood. This work underscores the necessity of using a suitable animal model to evaluate the potential risks associated with IVF and other ART in the reproductive systems of humans and mice. It also emphasizes the broad effects of IVF on various systems that have been overlooked and the importance of improving ART technologies to ensure the health of both offspring and future generations.

**Figure 8.**
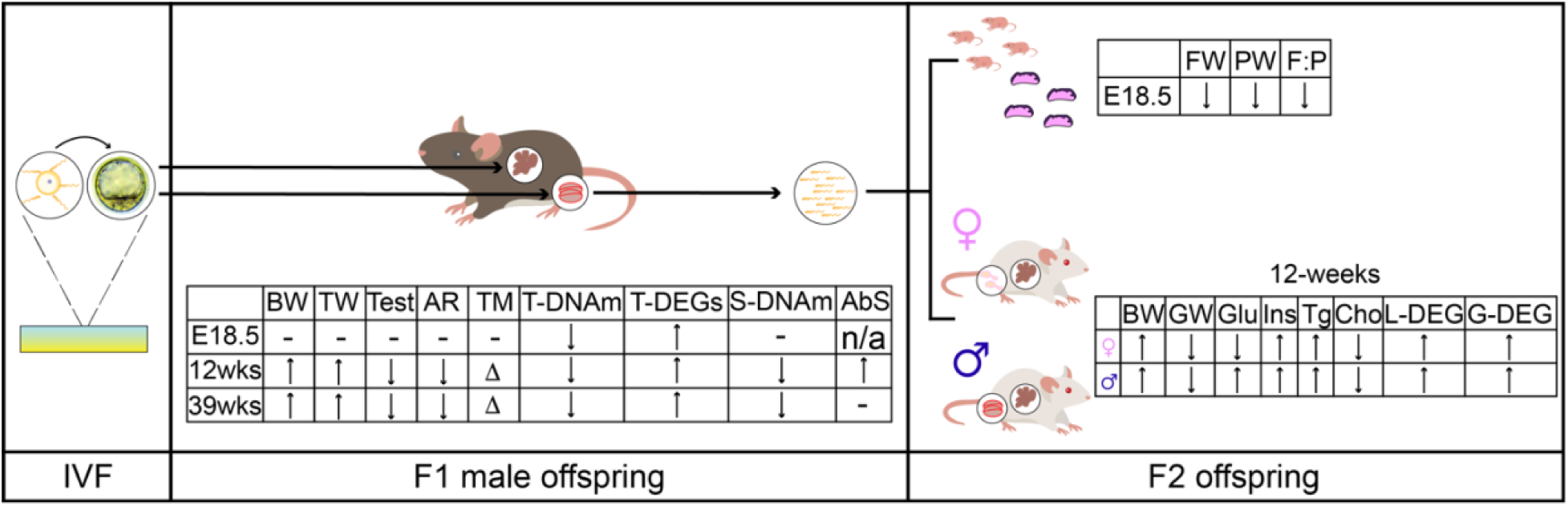
Summary of major findings. The use of IVF induced changes in testicular and sperm morphology and molecular profiles. Arrows indicate the direction of the changes compared to Naturals, with more arrows showing more significant changes. F1 male offspring: BW: Body weight, TW: Testicular weight, Test: Testosterone, AR: Androgen Receptor protein levels, TM: Testicular morphology, T-DNAm: Testicular DNA methylation, T-DEGs: Testicular differentially expressed genes, S-DNAm: sperm DNA methylation, AbS: Abnormal sperm morphology. F2 offspring: E18.5 FW: Fetal weight, PW: Placental weight; F:P: Fetal weight to placental weight ratio. 12-weeks: BW: body weight, GW: Gonad (testis or ovary) weight, Glu: Glucose, Ins: Insulin, Tg: Triglycerides, Cho: Cholesterol, L-DEGs: Liver differentially expressed genes, G-DEGs: Gonadal (testis or ovary) differentially expressed genes. ↑: significantly higher, ↓: significantly lower, Δ: changed, -: not measured in this project, n/a: not applicable.

## Methods

For this study, sex was considered as a biological variable in our experimental design, due to major differences in adverse outcomes previously observed (7, 11, 18). We included only males to ensure a comprehensive evaluation of the effects of IVF on the next generation. Our results cannot be generalized, as female and male reproductive development are different. Nevertheless, evaluation of the multigenerational impact of IVF includes both female and male offspring, as the second generation from an IVF male has not been previously studied, and no sex-specific effects have been identified.

### Animals

Breeding stocks of CF-1 female and male (Charles River, Wilmington, MA, USA), C57BL/6-Tg(CAG-EGFP)131Osb/LeySopJ male and SJL female (The Jackson Laboratory, Bar Harbor, ME) to generate SJL/B6 males as sperm donors, and CD-1 vasectomized male mice (Charles River, Indianapolis, IN) were maintained in a pathogen-free facility. All animals were housed in polysulfone cages and had access to drinking water and chow (Laboratory Autoclavable Rodent Diet 5010, LabDiet) *ad libitum*. All animal work was conducted with the approval of the Institutional Animal Care and Use Committee (IACUC) at the University of Pennsylvania. IACUC protocol 80354 has been previously revised and approved.

### Generation of Natural offspring

Offspring of conceived naturally (Naturals) were generated by mating CF-1 females in their natural estrous cycle with 8-week-old SJL/B6 males heterozygous for GFP. Detection of a vaginal plug marked embryonic day 0.5 (E0.5), and the embryos developed *in vivo* without embryo transfer (Figure 1A).

### Generation of in vitro fertilization offspring

As previously described (7, 11, 24, 25), IVF offspring were generated according to optimized protocols recommended by the Jackson Laboratory (56). CF-1 females were superovulated with 5 IU eCG followed by 5 IU hCG 46 h after eCG injection. On the day of IVF, sperm were collected from the vas deferens and caudal epididymis of a male SJL/B6 heterozygous for GFP. The collection was performed using EmbryoMax Human Tubal Fluid media (HTF, EMD Millipore) containing 3% w/v bovine serum albumin (BSA, AlbuMax, GIBCO) and under mineral oil (Irvine Scientific, USA). Sperm were then capacitated for at least 1 h. Eggs were collected and fertilized with capacitated SJL/B6 sperm in HTF medium. After 4 h, eggs with evident pronuclei (fertilized) were washed in HTF medium and EmbryoMax KSOM medium containing ½ amino acids (KSOM+AA, EMD Millipore) before culturing to the blastocyst stage in KSOM+AA covered with mineral oil (Irvine Scientific, USA) at 37°C in an atmosphere of 5% CO_2_, 5% O_2_, 90% N_2_. After 3.5 days of culture, blastocysts were removed from the KSOM+AA culture droplet and briefly washed in Multipurpose Handling Medium Complete (MHM-C, Irvine Scientific) with Gentamicin prior to embryo transfer. Pseudopregnant recipients (3.5 days post-copulation) were generated by mating CF-1 females with CD-1 vasectomized males. Each pseudopregnant recipient received ten blastocysts by non-surgical embryo transfer (Figure 1B). Day of blastocyst transfer was defined as E3.5.

### Fostering of natural and IVF offspring

Fostering of natural and IVF offspring was performed as previously described (11). We generated 3 cohorts using individuals from five different litters for each timepoint: I) E18.5 cohort of 12 Natural, and 10 IVF males; II) 12 weeks-of-age cohort of 12 Natural and 12 IVF males; and III) 39 weeks-of-age cohort of 12 Natural and 19 IVF males.

### Tissue collection

Naturally mated females or pseudopregnant recipients with IVF embryos were euthanized at E18.5, and fetuses and placentas were collected. Fetal and placental wet weights were recorded. Gonadal tissue was snap frozen in liquid nitrogen. Adults, at 12-weeks and 39 weeks-of-age were euthanized. Testes were collected and seminiferous tubules were collected from one of the testes and snap frozen in liquid nitrogen. The other testis was fixed in 10% of phosphate-buffered formalin for histological analyses. Sperm was collected from the vas deferens and cauda, processed using somatic cell lysis buffer, and the isolated sperm was snap frozen. Blood was collected, serum was then isolated by centrifugation at 4°C and kept at −80°C for hormone measurements.

### Sperm morphology analysis

Previously collected sperm from 12 weeks was used to obtain smears for morphological analysis using Hematoxylin-eosin staining. We assessed abnormal sperm, categorizing them as follows: abnormal head (%head), abnormal tail (%tail), abnormal mid-piece (%mid-piece), and total abnormal sperm (%abnormal). Our analysis referenced previous studies by other authors who also assessed sperm morphology (57, 58).

### Hematoxylin-eosin, Picrosirius red and Mason trichrome staining

Testicular sections were stained with hematoxylin-eosin as described (7, 24). Testicular sections were imaged using an EVOS FL Auto Cell Imaging System and software (Thermo Fisher Scientific, Waltham, MA, USA) at 4× magnification and analyzed using FIJI/ImageJ (National Institutes of Health). Changes in the seminiferous tubules area, thickness and overall structure were recorded. Image/J was used to identify changes in the intensity of the stain associated with collagen levels.

To assess changes in collagen deposition and potential fibrosis, picrosirius red (PR) staining was used on testicular sections as previously described (59). Briefly, tissue sections were deparaffinized in xylene and rehydrated through a graded series of ethanol concentrations (100%, 70%, and 30%), followed by rinsing with deionized water. Slides were then immersed in a PR staining solution, prepared by dissolving Sirius Red F3B (Direct Red 80, C.I. 35782, Sigma-Aldrich, St. Louis, MO, USA) in a 1.3% saturated aqueous picric acid solution (Sigma-Aldrich, St. Louis, MO, USA) at 0.1% w/v. After a 30-min incubation in the staining solution at room temperature, the slides were destained with 0.05 N HCl. The tissue sections were then dehydrated in 100% ethanol, cleared in xylene, and mounted with Permount mounting medium. Stained slides were imaged using an EVOS FL Auto Cell Imaging System (Thermo Fisher Scientific, Waltham, MA, USA). The percentage of PR-positive signal was calculated using three different testicular sections per animal with ImageJ.

To visualize collagen and validate PR staining, the Masson’s Trichrome (MT) assay was performed according to the manufacturer’s instructions (Polysciences, Inc., Warrington, PA, USA) on three testicular sections from each male. Slides were stained by the Pathology Core at the Children’s Hospital of Philadelphia (CHOP). Briefly, testicular sections were deparaffinized in xylene and rehydrated in graded ethanol (100% and 95%), followed by mordanting in preheated Bouin’s solution for 15 min at 60°C. Slides were then gently washed under running tap water for 5 min and stained with Weigert’s Iron Hematoxylin for 10 min. After several washes with tap water, the slides were stained in Biebrich Scarlet–Acid Fuchsin solution for 5 min and rinsed in distilled water. Samples were then incubated with phosphotungstic/phosphomolybdic acid for 10 min and finally stained with Aniline Blue for 5 min. After several rinses with distilled water, the slides were transferred to 1% acetic acid for 1 min, dehydrated in ethanol, cleared in xylene, and mounted in Vectashield mounting medium. Slides were imaged in brightfield using an EVOS FL Auto Cell Imaging System. Images were analyzed using FIJI/ImageJ and the plugin colour_deconvulation2 (60, 61). The percentage of MT-positive signal was calculated using three different testicular sections per animal.

### Testosterone levels

Serum collected from blood on the day of euthanasia was used to measure changes in testosterone levels using the commercially available Mouse/Rat Testosterone ELISA Kit (ab285350, Abcam, USA) and following the manufacturer’s guidelines.

### DNA and RNA isolation testis and sperm

DNA and RNA were isolated from one-quarter of each snap-frozen testis, as previously described (62). Sperm was collected from the vas deferens and caudal epididymis at 12 and 39 weeks-of-age and capacitated in EmbryoMax Human Tubal Fluid medium (HTF, EMD Millipore) with 3% w/v bovine serum albumin (BSA, AlbuMax, GIBCO) for 30 min at 37°C. Motile sperm was collected by removing the supernatant, spun down for 5 min at 650xg, and incubated for 15 min on ice with somatic cell lysis buffer (0.1% SDS, 0.5% Triton-X-100) to lyse and remove somatic cells. After treatment with the lysis buffer, sperm was counted, spun down for 5 min at 10,000xg, and snap-frozen for storage at −80°C until further processing. For DNA isolation, seminiferous tubules were digested in lysis buffer (50 mM Tris, pH 8.0, 100 mM EDTA, 0.5% SDS) with proteinase K (180 U/mL; Sigma-Aldrich) overnight at 55°C. Sperm pellets were resuspended in sperm lysis buffer (20 mM Tris-HCl, pH 8.0, 200 mM NaCl, 20 mM EDTA, 4% SDS) with 5 μL of β-mercaptoethanol and proteinase K (180 U/mL) and incubated overnight at 55°C. Genomic DNA was isolated using phenol:chloroform:alcohol (25:24:1; Sigma-Aldrich), followed by ethanol precipitation and resuspension in TE buffer (10 mM Tris-HCl, pH 8.0, 0.5 mM EDTA).

RNA was isolated from a cross-section of the testis using an adapted protocol of trizol (Thermo Fisher Scientific) and Qiagen Micro RNA Kit (Qiagen, Germany), following the manufacturer’s protocol. DNAse treatment was performed during RNA isolation to eliminate genomic DNA contamination. RNA quality and concentration were determined by RNA ScreenTape analysis using a TapeStation (Agilent Technologies, USA).

### Luminometric methylation assay

Genomic DNA (1 µg) from testis, sperm and ovaries (second generation) was used to measure global DNA methylation by luminometric methylation assay as previously described (7, 24, 25).

### Genome-wide DNA methylation profiling and analysis

We performed genome-wide DNA methylation profiling on a random subset of 12-week and 39-week-old offspring sperm (n=6 males for each group, 12 for each age), as previously described (7, 35). The samples were processed at the Center for Applied Genomics Genotyping Core at the Children’s Hospital of Philadelphia. Bisulfite-treated DNA (1 µg) was pipetted onto an Illumina Infinium Mouse Methylation-12v1-0 BeadChip (Illumina, CA, United States) that was run on an Illumina iScan System (Illumina, CA, United States) using the manufacturer’s standard protocol as previously described (5, 35, 36).

Raw IDAT files were processed, as previously described (5, 35, 63). Changes in DNA methylation were identified by comparing the methylation levels probe by probe between Naturals and IVF. We analyzed the data using the pipeline recommended by the SeSAMe package (64), as previously described (7, 35). CG identity was assessed using the tool KYCG part of the SeSAME R package. Pathway analyses were performed using the package for R: clusterProfiler. The raw methylation array data reported in this work has been deposited in the NCBI Gene Expression Omnibus under accession number GSE280286.

### RNA sequencing

We performed RNA sequencing on a random subset of E18.5, 12 and 39-week testis from n=5 individuals for each group, different litters and tissue. We also profiled liver and ovaries of the second generation. As previously described (7, 11), total RNA (4 µg) was used to prepare mRNA-sequencing libraries using KAPA mRNA-Seq library synthesis kit and KAPA Single-Indexed adapter kit (Kapa Biosystems, Wilmington, MA, USA). Library quality control was conducted using High Sensitivity DNA ScreenTape for TapeStation (Agilent Technologies, USA) and Kapa Library Quantification Kit (Kapa Biosystems, Wilmington, MA, USA). Sequencing was performed using the NovaSeq 1000 platform (Illumina, San Diego, CA).

RNA-seq reads were analyzed as previously described (7, 11). Numbers for total sequenced, aligned, and counted reads for each sample are listed in Supplemental Table 1. The raw sequencing data reported in this work has been deposited in the NCBI Gene Expression Omnibus under accession number GSE280286.

### Real-time PCR

Extracted testicular RNA from Naturals and IVF offspring at 12 and 39 weeks were cDNA converted as previously described (5, 7, 11). Primers are listed in Supplemental Table 2. Relative expression was calculated using the quantified expression from the endogenous control *Nono* and Actin B (*ActB*) that showed stable expression levels in mouse across multiple samples. RT-PCR was conducted on all samples, including those used for RNA-seq: 12-weeks Natural (n = 12), IVF (n = 12); and 39-week Naturals (n = 12) and IVF (n = 12).

### Western Blot

Seminiferous tubules from 12- and 39-week males were used for protein analysis by Western Blot as previously described (11). Protein lysate (20 μg) was used per sample and each membrane was probed with primary antibodies diluted in 5% nonfat dairy milk in TBS-T for GAPDH (1:5,000, Cell Signalling) and either AR (1:1,000; Abcam, Cambridge, UK), VCAM1 (1:1,000; Abcam, Waltham, MA, USA), or COL1A1 (1:1,000; Abcam, Waltham, MA, USA) Levels of AR, COL1A1 and VCAM1 were determined relative to GAPDH levels and compared between groups.

### Multigenerational studies in natural and IVF offspring

12-week-old Natural and IVF offspring were mated with wild-type CF-1 females for four consecutive litters. At E18.5 some pregnant females were euthanized, and both fetus and placenta were collected to assess prevalence of the phenotype on the next generation. Other litters were weaned at 21 days-of-age, and using a calibrated digital scale, body weights were measured at birth and weekly from 3-12-weeks-of-age. Five animals were housed per cage to ensure no differences in food and water consumption as previously described (7, 11). Twelve-week-old animals were euthanized, and blood glucose levels were obtained by tail snip using a hand-held glucometer (ReliOn, Bentonville, Arkansas, USA). Whole blood was collected by cardiac puncture, centrifuged, and serum was collected and stored at −80°C for subsequent assays. Total triglycerides were assayed using enzymatic colorimetric assay kits from Stanbio (Boerne, TX, USA) and insulin was assayed using enzymatic colorimetric assay kits from Crystal Chem (Illinois, USA). Total cholesterol, HDL, and LDL/VLDL were assayed using the Abcam HDL and LDL/VLDL Cholesterol Assay Kit (ab65390; Cambridge, UK).

Organ weights were recorded, and blood was collected for metabolic and hormonal panels. For molecular analysis, part of the liver, previously homogenized pancreas in TRIzol, one of the ovaries, and one of the testes were snap-frozen in liquid nitrogen were used. For morphological and histological analysis, the remaining liver, pancreas, one of the ovaries, and one of the testes were fixed in 10% formaldehyde in PBS.

### Statistical analyses

All samples were statistically analyzed an unpaired t-test. Probability of genomic regions in the array were compared using a Bernoulli distribution using R v 4.2.2 (R foundation for Statistical Computing; www.R-project.org/). Significant differences comparing Natural and IVF offspring were denoted with as statistically significant if *P*<0.05. Correlation analysis was performed using a Pearson r test and consider significant if *P*<0.05. All statistical analyses were performed using GraphPad Prism version 9.1.

### Data availability

All values for all data points in graphs are reported in the Supporting Data Values file. Additionally, the data array and sequencing data underlying this article is available at NCBI Gene Expression Omnibus at https://www.ncbi.nlm.nih.gov/geo/, under GEO accession number GSE280286.

## Author Contributions

EAR-C and MSB designed research studies. EAR-C, AS, CNH and LN conducted experiments. EAR-C and AS acquired data. EAR-C wrote the manuscript; and all authors edited and reviewed the manuscript prior to submission. ER-C is listed as first author based on his conceptualization and initiation of the project.

## Acknowledgements

We acknowledge the individuals that helped to make this work possible, including Chris Krapp, for his technical expertise; Ken Zaret for use of microtome; CHOP Pathology Core Laboratory for use of embedding equipment, Penn Cell and Developmental Biology Microscopy core for use of the imaging facilities and the Center for Applied Genomics Genotyping Core at CHOP for performing the Infinium Mouse Methylation BeadChip assays. This work was funded by a National Centers for Translational Research in Reproduction and Infertility grant HD068157 (MSB), Ruth L. Kirschstein National Service Award Individual Postdoctoral Fellowship HD107914 (ER-C) and National Institute of Health Training program in Cell and Molecular Biology T32 GM007229 (CH).

