## Supplemental Figures for "*In Vitro* Fertilization induces reproductive changes in male mouse offspring and has multigenerational effects"

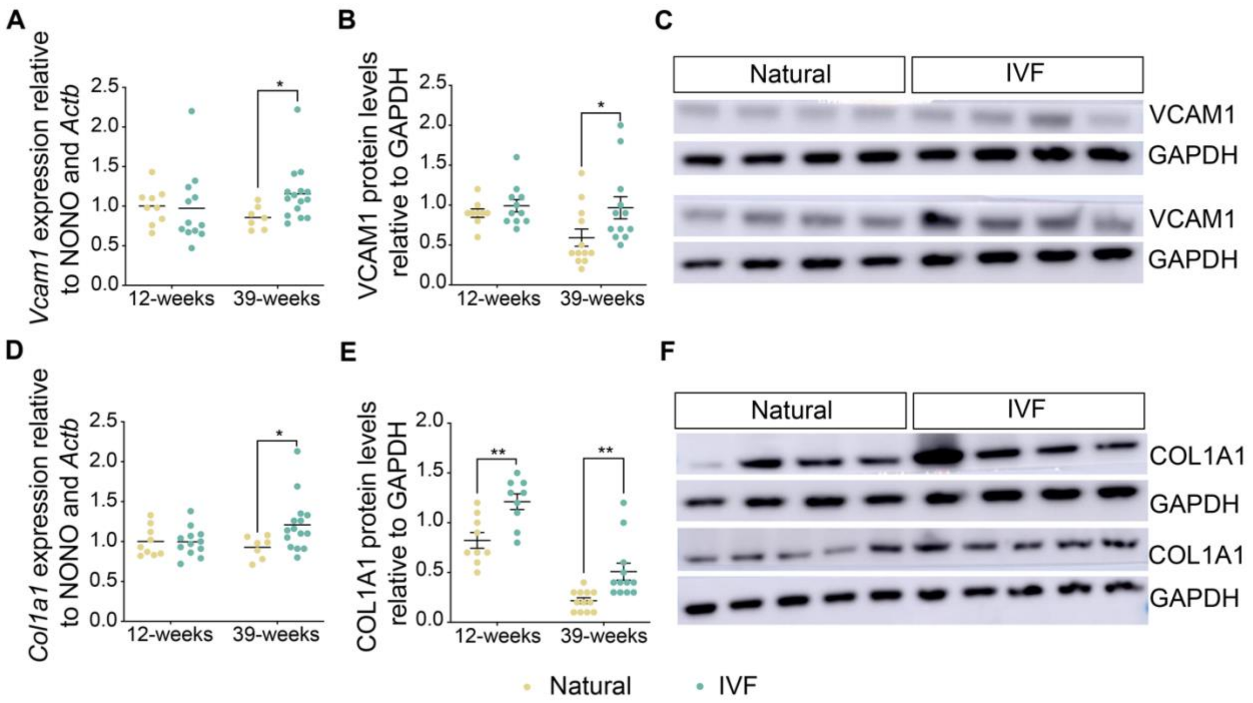


**Supplemental Figure 1 (corresponding to main Figure 2).** Gene expression and protein levels of *Col1a1* and *Vcam1*. Real time PCR using RNA from testis from 12- and 39-week-old offspring to measure expression of (A) *Vcam1*, and (D) *Col1a1* relative to *Nono* and *Actb*. Western blot for protein in whole testis for (B) VCAM1 and (E) COL1A1are relative to GAPDH, (C) and (F) western blots images corresponding to each protein. Data are depicted as mean±s.e.m, *n*=10-15 per group. The black line represents the mean of each group. Statistical significance was determined by t- test, *P<0.05, and **P<0.01 when compared groups against Natural.

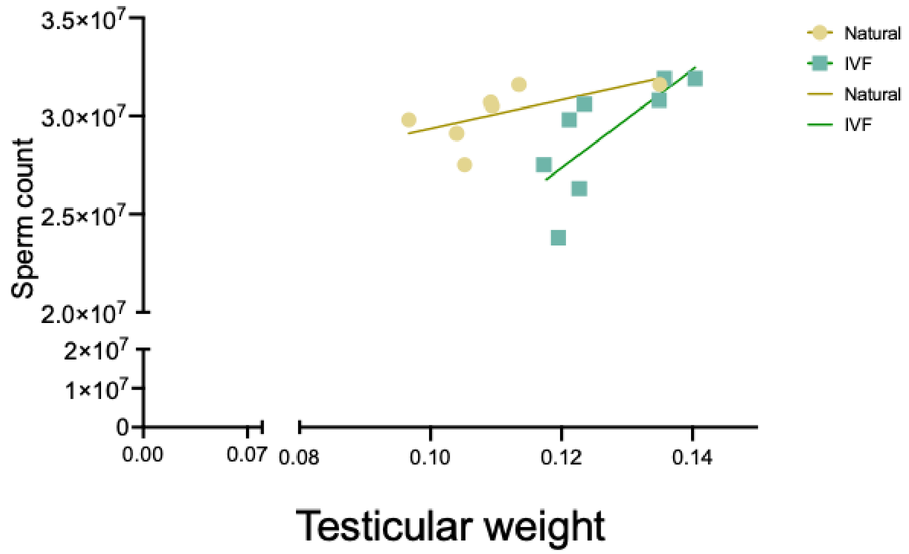


**Supplemental Figure 2 (corresponding to main Figure 3).** Correlation between sperm count and testicular weight in both IVF and Natural offspring.

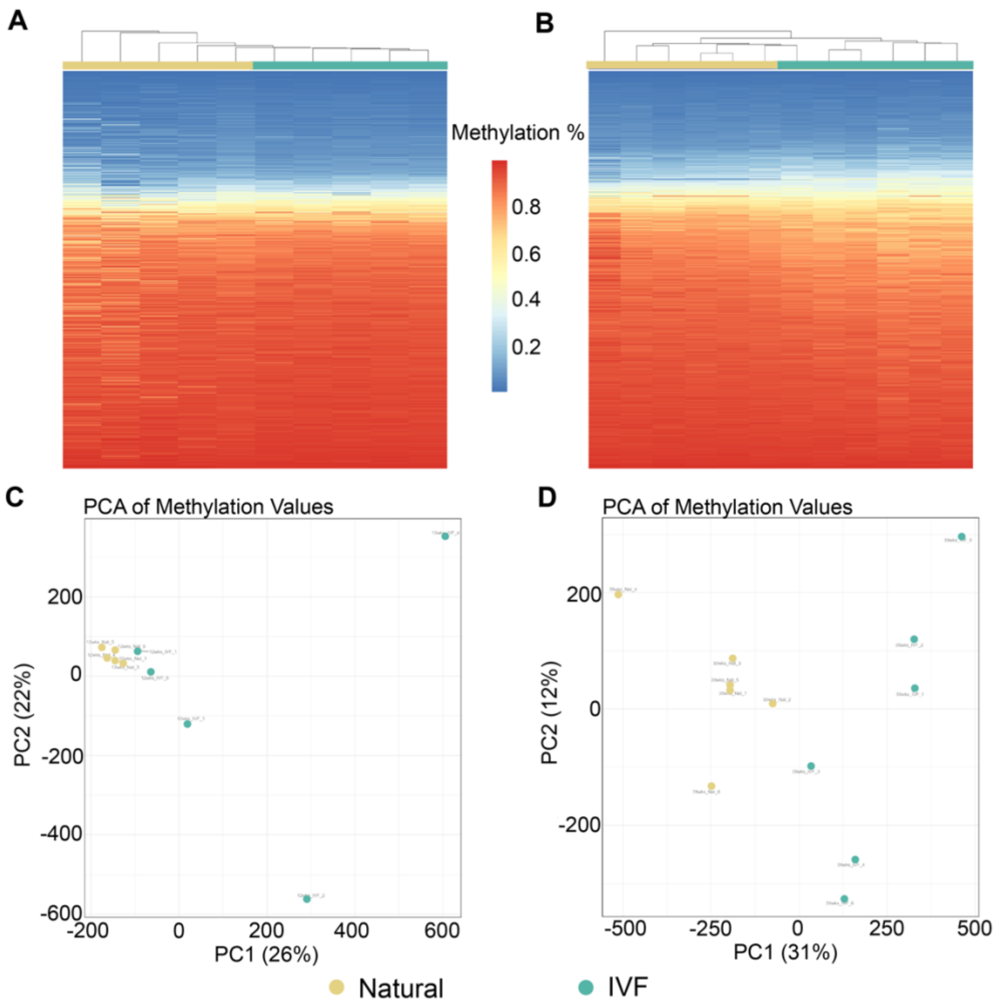


**Supplemental Figure 2 (corresponding to main Figure 3).** Heatmap of all the probes from the Illumina Beadchip array using sperm: (A) 12-weeks-of-age, and (B) 39-weeks-of-age. Principal component analysis (PCA) for sperm samples run in the Illumina Beadchip array: (C) 12-weeks-of-age, and (D) 39-weeks-of-age.

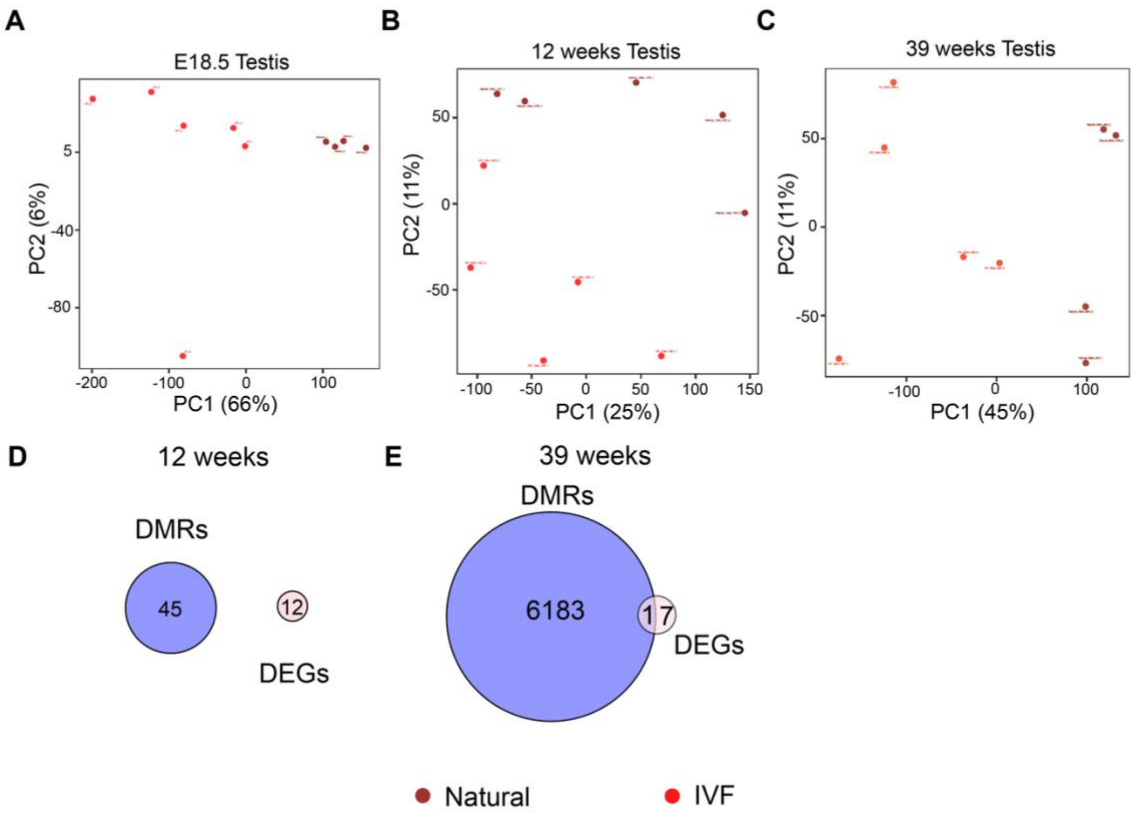


**Supplemental Figure 3 (corresponding to main Figure 4).** Principal component analysis (PCA) for RNAseq results. PCA before DEG analysis for (A) E18.5 Testis, (B) 12 weeks Testis, (C) 39 weeks Testis. Overlap between sperm DMRs and DEGs testis: (D) 12 weeks Testis and (E) 39 weeks Testis.

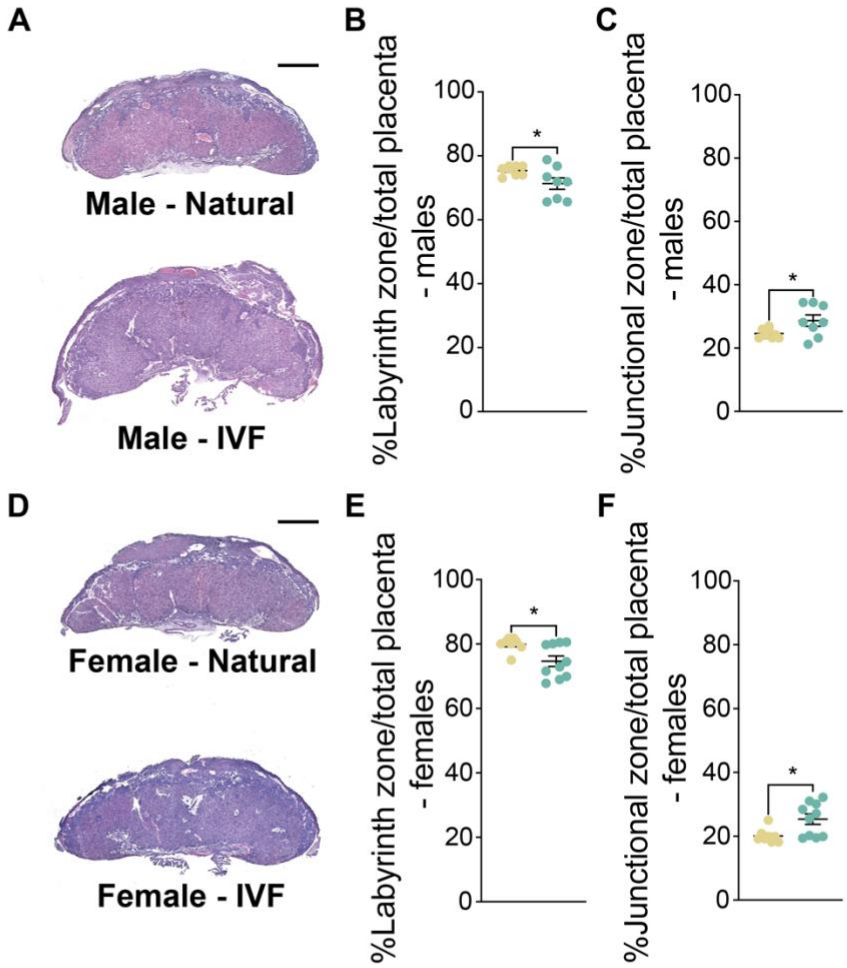


**Supplemental Figure 4 (corresponding to main Figure 5).** Percentage of placental junctional and labyrinth zone at E18.5 using hematoxylin-eosin staining. Placenta cross-sections from (A) hematoxylin-eosin histological cuts from E18.5 F2 male placentas with the percentage of (B) labyrinth zone and (C) junctional zone shown. (D) hematoxylin-eosin histological cuts from E18.5 F2 female placentas with the percentage of (E) labyrinth zone and (F) junctional zone indicated. Scale bar: 850 µm. Each data point represents an individual conceptus from a minimum of four different litters. The black line represents the mean of each group (n=8/group/sex). Statistical significance was determined by t- test, *P<0.05 when compared groups against Natural.

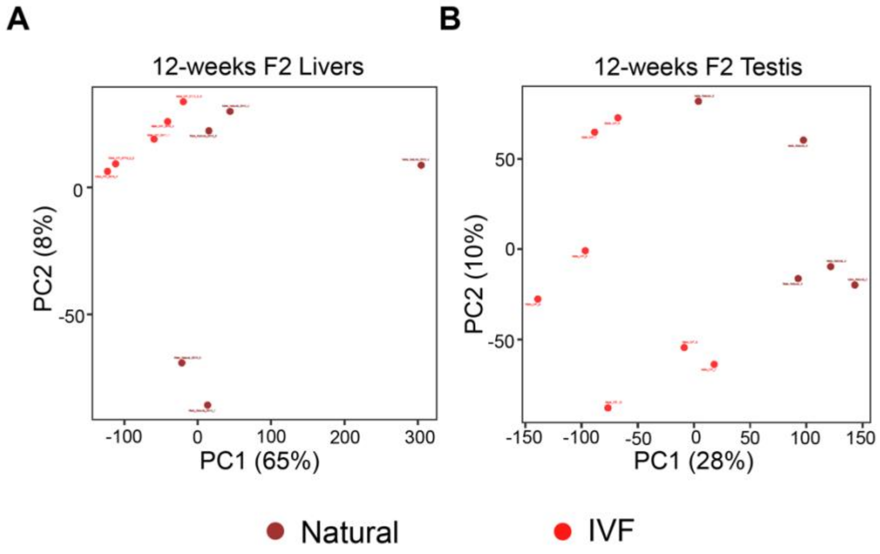


**Supplemental Figure 5 (corresponding to main Figure 6).** Principal component analysis (PCA) for RNAseq results. PCA before DEG analysis for male second generation (A) livers, (B) testis.

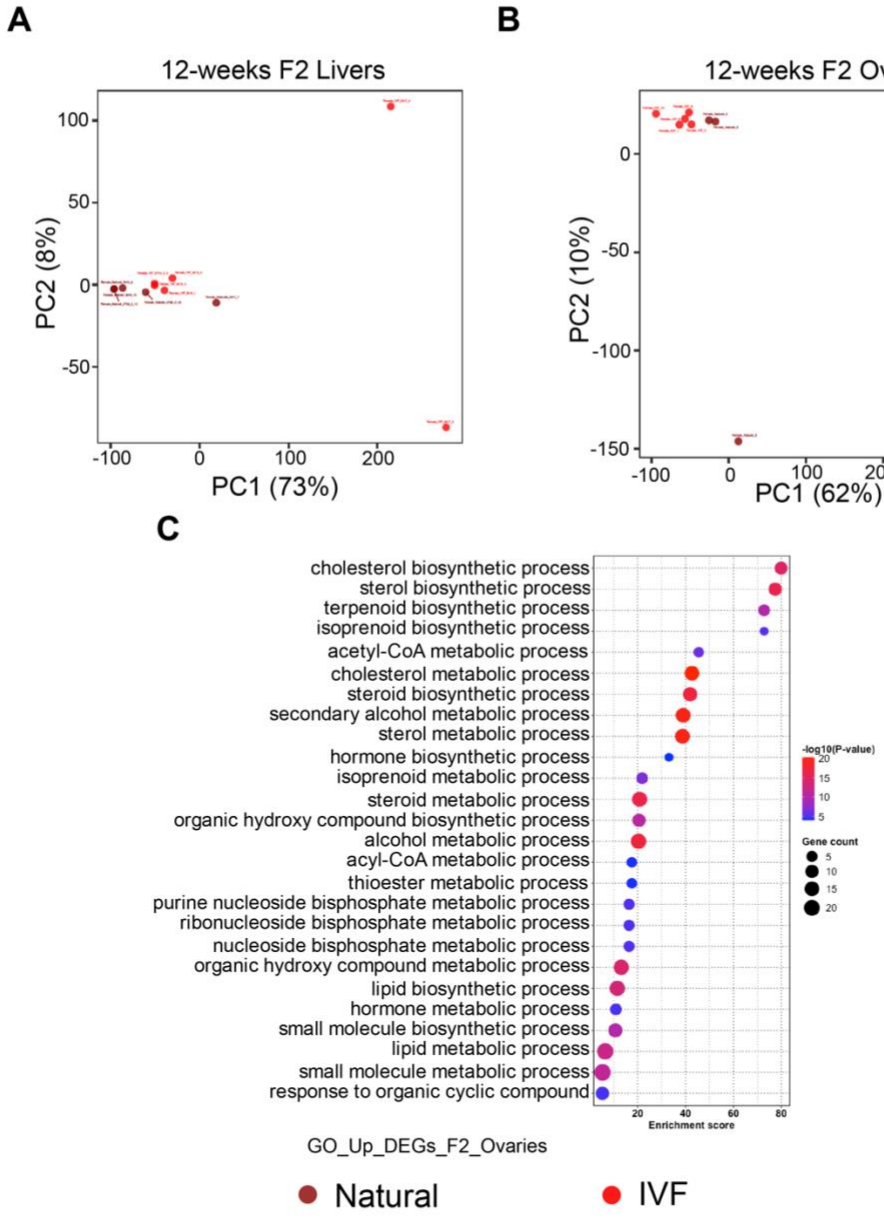


**Supplemental Figure 6 (corresponding to main Figure 7).** Principal component analysis (PCA) for RNAseq results. PCA before DEG analysis for female second generation (A) livers, (B) ovaries. (C) Gene ontology analysis for upregulated genes in ovaries from second generation females.

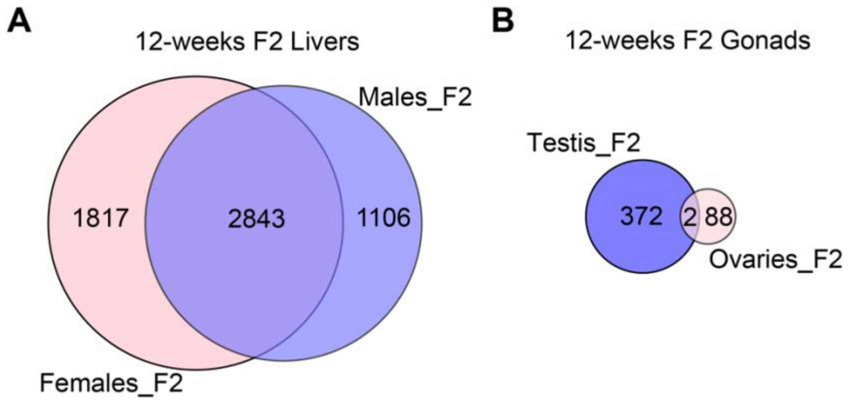


**Supplemental Figure 7 (corresponding to main Figure 6 and 7).** Venn diagrams showing common DEGs males and females for (A) Livers and (B) Gonads.

Supplemental Table XXX: Numbers for total sequenced, aligned, and counted reads for each sample RNA seq.

Supplemental Table XXX: Real time PCR primers
